# The Southern Bluefin Tuna mucosal microbiome is influenced by husbandry method, net pen location, and anti-parasite treatment

**DOI:** 10.1101/2020.05.19.105270

**Authors:** Jeremiah J Minich, Cecilia Power, Michaela Melanson, Rob Knight, Claire Webber, Kirsten Rough, Nathan J. Bott, Barbara Nowak, Eric E Allen

## Abstract

Aquaculture is the fastest growing primary industry worldwide. Marine finfish culture in open ocean net pens, or pontoons, is one of the largest growth areas and is currently the only way to rear high value fish such as bluefin tuna. Ranching involves catching wild juveniles, stocking in floating net pens and fattening for four to eight months. Tuna experience several parasite-induced disease challenges in culture that can be mitigated by application of praziquantel (PZQ) as a therapeutic. In this study, we characterized the microbiome of ranched southern Bluefin Tuna, *Thunnus maccoyii*, across four anatomic sites (gill, skin, digesta, and anterior kidney) and evaluated environmental and pathological factors that influence microbiome composition, including the impact of PZQ treatment on microbiome stability. Southern bluefin tuna gill, skin, and digesta microbiome communities are unique and potentially influenced by husbandry practices, location of pontoon growout pens, and treatment with the antiparasitic PZQ. There was no significant relationship between the fish mucosal microbiome and incidence or abundance of adult blood fluke in the heart or fluke egg density in the gill. An enhanced understanding of microbiome diversity and function in high-value farmed fish species such as bluefin tuna is needed to optimize fish health and improve aquaculture yield. Comparison of the bluefin tuna microbiome to other fish species, including *Seriola lalandi* (yellowtail kingfish), a common farmed species from Australia, and *Scomber japonicus* (Pacific mackerel), a wild caught Scombrid relative of tuna, showed the two Scombrids had more similar microbial communities compared to other families. The finding that mucosal microbial communities are more similar in phylogenetically related fish species exposes an opportunity to develop mackerel as a model for tuna microbiome and parasite research.

## Introduction

Bluefin tuna is one of the highest value fish in the world. In 2014, the three species of bluefin including Atlantic Bluefin Tuna (ABT), Pacific Bluefin Tuna (PBT), and Southern Bluefin Tuna (SBT) had a combined total value of USD$610-660 million for fishers with a total value of USD$2-2.5 billion at final sale (Macfadyen, 2016). To meet high demands, bluefin tunas are ranched in open ocean net pens, or pontoons, which currently represents approximately 20% of total global bluefin production. In Australia, the SBT wild-caught fishery began in 1949 (Serventy, 1956), peaking in 1982 at an annual catch of 21,000 tons (Geen and Nayar, 1989); after this peak catch restrictions were imposed on the fishery. Ranching of SBT involves the collection and transfer of wild juvenile tuna into static pontoon enclosures where they are reared and fattened for up to eight months. Over this period tuna are fed whole baitfish, which results in a near doubling of fish biomass. Ranching began in Port Lincoln Australia in 1990 and in 2017-18 had an annual production value of AUD$166 million (EconSearch; François et al., 2010; Ellis and Kiessling, 2016). One of the primary challenges to promoting and ensuring long term success of the industry is to maintain and improve fish health.

When cultured in open ocean pontoons, fish are exposed to free-living microbes, including opportunistic pathogens and infectious parasites (Nowak, 2007). Several parasites have been identified in ranched SBT including *Miamiensis avidus* (scuticociliate), *Caligus* sp. (copepod ‘sea lice’), *Cardicola* spp. (blood flukes), and *Hexostoma thynni* (gill fluke) (Nowak et al., 2003). Digenean blood flukes, specifically *Cardicola* spp., infect all three species of bluefin tuna during ranching which can lead to morbidities and mortalities causing significant losses for producers (Balli et al., 2016; Ellis and Kiessling, 2016). Among the most destructive parasites for SBT are blood flukes that include two species, *Cardicola fosteri* and *C. orientalis* (Cribb et al., 2000; Colquitt et al., 2001). In captivity, infections peak approximately two months post-relocation to pontoons with SBT having on average 27 flukes per fish (Aiken et al., 2006). Blood flukes produce eggs which are deposited in the gills (up to 9 million eggs per individual) resulting in morbidity and mortality (Colquitt et al., 2001; Bullard and Overstreet, 2002; Shirakashi et al., 2012b). Currently, praziquantel (PZQ) is the only therapeutic used to treat *C. fosteri* and *C. orientalis* infections. PZQ is administered orally through feeds by injecting 75-150 mg/kg body weight into sardines which are then fed to SBT (Hardy-Smith et al., 2012). Treatment with PZQ (15 mg/kg body weight) has a positive impact on adults SBT health by eradicating adult flukes and has shown to be effective in PBT juveniles as well (Shirakashi et al., 2012a). Lower doses (3.75-7.5 mg/kg) have also shown positive effects for adult fluke eradication with organ elimination after 24 hours (Ishimaru et al., 2013). In addition to therapeutic treatments, pontoon location and alterations to husbandry practices, such as ranching at greater depths, can also lower infection outcomes with a combined impact of decreasing mortalities from 15% to <1% (Kirchhoff et al., 2011).

Marine fish harbor specialized microbiomes within the gill, skin, and gastrointestinal tract that are distinct from the surrounding seawater and influenced by environmental variables (Minich et al., 2019a). These microbial communities play important roles in mucosal barrier defenses to protect against pathogen invasion and maintain host health. Manipulation and optimization of the microbiome of farmed fish through prebiotic, probiotic, or synbiotic interventions are active areas of research to promote fish health and resilience (Gomez et al., 2013; Kelly and Salinas, 2017). Understanding how the fish mucosal microbiome is impacted by the environment and pathological status of the host, e.g. parasite infection load, is largely unknown yet may be important for the development of therapeutics and/or diagnostics (Li et al., 2016). Secondary infections with bacteria or viruses can also be enhanced during parasitic infections but little is known about blood flukes specifically (Kumon et al., 2002; Boxaspen, 2006; Lhorente et al., 2014; Novak et al., 2016). Since blood fluke egg deposition in the gills leads to pathology and respiratory problems, evaluating microbiome changes between infected and non-infected individuals may reveal dysbiosis signatures that further influence host health. Moreover, the presence of a parasite infection itself may lead to changes in the mucosal bacteria communities of the fish which could, in turn, be used to predict, diagnose, and monitor parasite infections. In Atlantic salmon, skin infections by sea lice resulted in decreased richness of skin bacteria which was further driven by an enrichment of potential pathogenic microbes including *Vibrio* spp., *Pseudomonas* spp., and *Tenacibaculum* spp. (Llewellyn et al., 2017). Understanding the mechanisms and interactions between infection, treatment, host response, and the microbiome, the collective “pathobiome” (Bass et al., 2019), are thus important for reducing the impact of infections.

In this study, ranched SBT were sampled to characterize the microbial diversity associated with mucosal body sites, including gill, skin, and gut, providing the first assessment of microbiome diversity in this ecologically and commercially important fish species. PZQ treatment has previously been shown to reduce *Cardicola* spp. parasite egg abundances in the gills (Power et al., 2019). To better establish the impact of antiparasitic treatment on microbiome composition, we analyzed mucosal microbiome composition as a function of parasite infection and PZQ treatment. We hypothesized that prevalence of *Cardicola* eggs in the gill would correspond to an altered microbial community. In addition, we hypothesized that among the body sites evaluated, the gastrointestinal tract would have the highest probability of being impacted by the oral treatment of PZQ delivered through feeds. During the course of a harvest event, a total of 65 SBT across six different pontoons from five spatially separate companies were sampled. Assessment of blood fluke prevalence, gill egg densities, and fish biometrics (length and weight) were collected and analyzed with respect to mucosal microbiome composition. Lastly, we compare the mucosal microbiome of SBT to another Scombrid species to evaluate its utility as a possible model organism for marine fish microbiome studies. A meta-analysis of the SBT microbiome compared to two other commercially important fish species suggests the Pacific chub mackerel, *Scomber japonicus*, may be a suitable model to study microbiome dynamics in marine aquaculture species, including bluefin tuna.

## Materials and Methods

### Experimental animals and sampling

Southern Bluefin Tuna ‘SBT’, *Thunnus maccoyii*, reared in ocean pontoons (also known as ranching) in Port Lincoln, Australia, were opportunistically sampled during a harvest event from July 10 - 20 2018. A total of 65 fish were sampled across six pens from five total companies. Between 9 and 12 fish were sampled from each pen. Fish were measured for total length (mm) and mass (grams) and Fulton’s condition factor ‘K-factor’ was calculated (Froese; FULTON, 1904). Fish (n=45) from four of the six pens were treated with the antiparasitic drug, anthelmintic praziquantel (PZQ) orally (injected into baitfish) at a dose of 30 mg/kg bodyweight 4-5 weeks after transfer to pontoons in a single treatment over two consecutive days by the prescribing veterinarian (APVMA Permit Number-Per 85738) (Power et al., 2019) while fish in the other two pens did not get treated with PZQ (n=20). Presence of adult flukes in heart and egg densities within the gill were measured per fish as previously described (Aiken et al., 2006; Power et al., 2019). Comparison of the SBT microbiome to other fish samples, including Pacific chub mackerel (MKL) and yellowtail kingfish (YTK), was performed computationally within Qiita (see below for data availability). All MKL samples were collected from the wild while the YTK were farmed. Details for fish sample collections are listed in the meta-analysis section below.

### Microbiome assessment

SBT microbiome samples were collected from four body sites (gill, skin, digesta, and anterior kidney) from each of the 65 fish for a total of 260 samples. Gill and skin communities were collected by swabbing approximately 25 cm^2^ surface area using cotton swabs. For digesta and anterior kidney samples, approximately 100 mg of fecal matter or tissue was collected and placed in a 1.5 ml tube. To preserve microbiome integrity, all microbiome samples were stored in 95% ethanol for 4-6 hours at room temperature while on the boat and then later transferred to a −20°C freezer where samples were stored until DNA extraction (approximately 2 weeks later) (Song et al., 2016).

Molecular methods outlined in the Earth Microbiome Project (earthmicrobiome.org) were used to process samples. Specifically, DNA was extracted in single tubes to avoid well-to-well contamination (Minich et al., 2019b) using the MoBio PowerSoil kit. Multiple titration replicates of 10 fold serial dilutions of positive controls (n=21) (*Escherichia coli*) were included to enable detection of background contaminates and determine the limit of detection of the method using the Katharoseq method (Minich et al., 2018b) (Minich et al., 2020). Miniaturized (5ul) PCR reactions were used to amplify genomic DNA (200 nl) (Minich et al., 2018a) using 16S rRNA V4 515/806 EMP primers (Parada et al., 2016; Walters et al., 2016). Equal volumes of amplicons (1 ul) were pooled across samples and processed through the Qiagen PCR cleanup kit and then sequenced on a MiSeq (Caporaso et al., 2012). Sequencing analysis was performed using Qiime2 (Bolyen et al., 2019, 2) and Qiita (Gonzalez et al., 2018).

### Statistics and meta-analysis

Samples were trimmed to 150 bp and then demultiplexed and processed through the deblur (Amir et al., 2017) pipeline to generate unique sOTUs (sub-Operational Taxonomic Units) also referred to as ASVs (Amplified Sequence Variants). Samples were rarified to 1000 reads as determined by Katharoseq cutoff. All downstream analyses were performed using the rarified biom table. Two sOTUs were removed from the dataset. The first was the sample used as a positive control (Enterobacteriaceae: TACGGAGGGTGCAAGCGTTAATCGGAATTACTGGGCGTAAAGCGCACGCAGGCGGTTTGTTAAGTCAGATGTGAAATCCCCGGGCTCAACCTGGGAACTGCATCTGATACTGGCAAGCTTGAGTCTCGTAGAGGGGGGTAGAATTCCAGG) and the second was a known PCR mastermix contaminant (Eisenhofer et al., 2018) (*Pseudomonas veronii*: TACAGAGGGTGCAAGCGTTAATCGGAATTACTGGGCGTAAAGCGCGCGTAGGTGGTTTGTTAAGTTGGATGTGAAATCCCCGGGCTCAACCTGGGAACTGCATTCAAAACTGACTGACTAGAGTATGGTAGAGGGTGGTGGAATTTCCTG) which we observe disproportionally in low biomass positive controls. Alpha diversity was calculated using richness (total unique sOTUs), Shannon evenness, and Faith’s Phylogenetic diversity. Alpha diversity comparisons were calculated using non-parametric Kruskal-Wallis (Kruskal and Wallis, 1952) test with multiple comparisons done using Benjamini-Hochberg with a 0.05 (Benjamini and Hochberg, 1995). Pairwise comparisons were performed using a non-parametric Mann-Whitney test (Mann and Whitney, 1947). Beta diversity was calculated using both weighted and unweighted UniFrac distances (Lozupone and Knight, 2005; Lozupone et al., 2010). To test which metadata categories or variables were associated with beta diversity, multivariate statistical testing was done using ADONIS which is a modified version of PERMANOVA (Anderson, 2001). Specifically, the following categorical variables were assessed for their impact on the microbiome: sample type, company, and PZQ treatment. Following those results, body sites were independently assessed for the impacts of PZQ treatment, company, parasite eggs in gill, parasite flukes in heart, tuna condition factor, tuna length, and tuna mass.

A metanalysis was performed to test the hypothesis that phylogenetically similar fish species have more similar microbial communities. Both alpha and beta diversity were compared. To do this analysis, publicly available microbiome data from gill, skin, and digesta microbiomes of Pacific chub mackerel (MKL), *Scomber japonicus,* (Qiita ID 11721, ERP2664334) and yellowtail kingfish (YTK), *Seriola lalandi,* were combined with SBT using the Qiita database. Mackerel samples were wild fish sampled throughout 2017 caught off the Scripps Institution of Oceanography Pier in the Eastern Pacific Ocean, San Diego. The yellowtail kingfish was sampled from pontoons in the Western Pacific Ocean, NSW Australia. Mackerel are phylogenetically closer to SBT in the Scrombridae family (mackerels, bonitos, and tunas) (Collette, 1983; Collette et al., 2001), whereas yellowtail are within the Carangidae family (jacks and pompanos) (Swart et al., 2015). For specimens collected in this study, yellowtail are however more similar to SBT in terms of trophic level, geographic location of sampling, and farmed rather than wild caught.

## Results

### Sampling design and results

Southern Bluefin Tuna were sampled during annual harvest across 10 days (July 10 2018 to July 20 2018) in Port Lincoln, Australia. A total of six pens were sampled among five different companies (Figure 1a). Two of the pens (n=20 fish) were not treated with the anthelmintic praziquantel (PZQ) while the other four pens (n=45 fish) were treated. Four anatomical sites were sampled for microbiome analysis (Figure 1b) including gill, skin, digesta, and anterior kidney. Fish biometrics including fork length (mm), mass (grams), and condition factor ‘K-factor’ were measured for 62 fish whereas data from three fish from July 15^th^ (host_subject_ID: SBT_51, SBT_57, and SBT_58) were not recorded (Figure 1). Tuna ranged in length from 920 mm to 1180 mm with a median length of 1070 mm (Figure 1c). Tuna ranged in mass from 16200 grams to 35000 grams with a median mass of 25950 grams (Figure 1d). The K-factor ranged from 1.81 to 2.73 with a median of 2.10. Blood fluke counts and parasite egg counts in the gill were recorded for all 65 fish. Blood fluke counts ranged from 0 (30 fish) to 6 with a median and mean intensity of 1 (Figure 1f). Parasite egg counts in the gill ranged from 0 (4 out of 65) to 3.94, with a median of 0.45 and mean intensity of 0.846 eggs per mm filament (Figure 1g).

**Figure 1.**
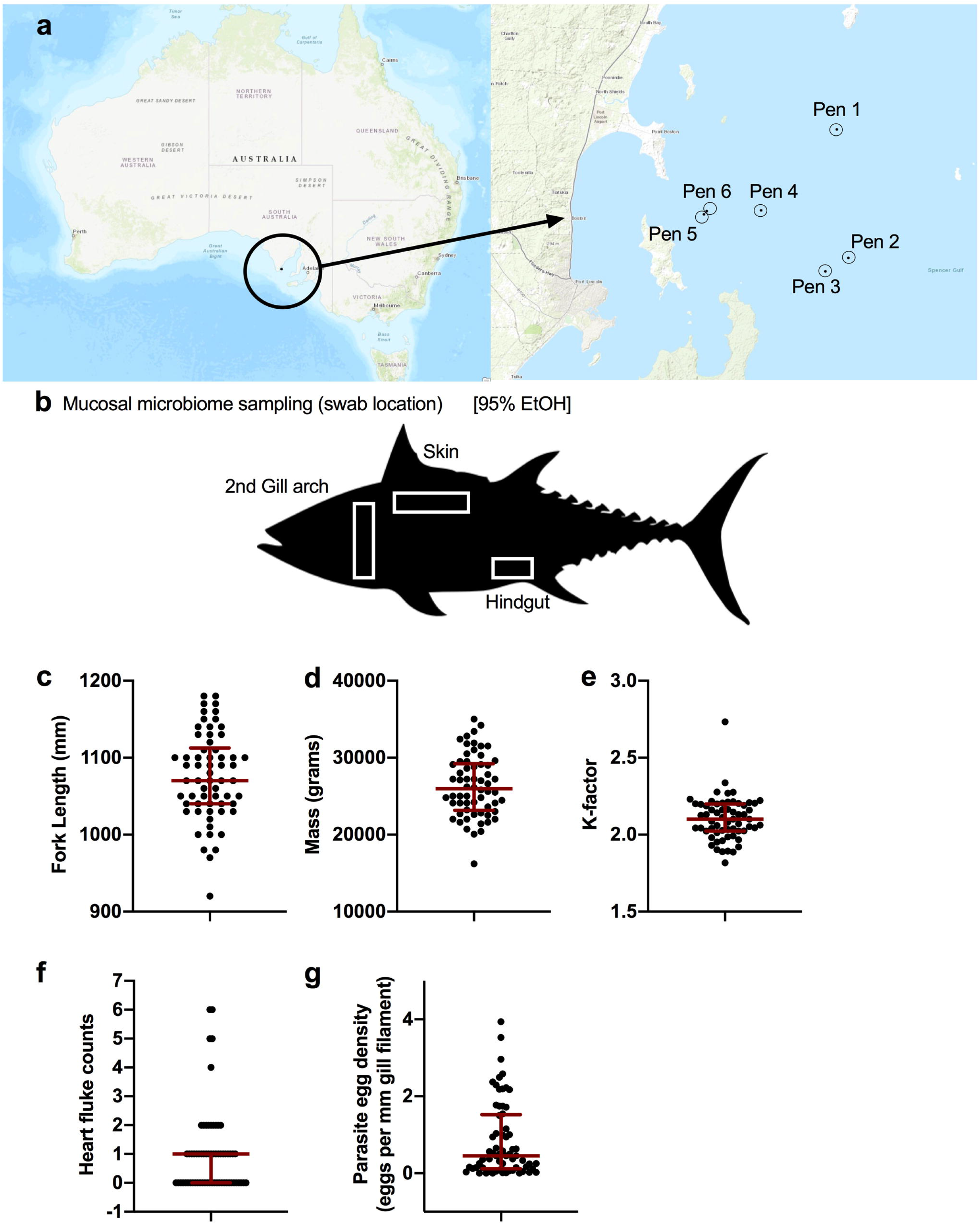
Sampling design of ranched SBT from Port Lincoln, Australia. a) Fish were sampled from a total of six pens spanning five companies. b) Upon harvest, four body sites were sampled and stored in 95% EtOH for the microbiome including the gill, skin, digesta (hindgut), and anterior kidney. Fish biometrics across all samples collected for c) fork length (mm: median, IQR), d) mass (grams: median, IQR), and e) calculated condition factor (median, IQR). Blood fluke parasite counts from f) heart (flukes per heart: median, IQR), and g) gill (eggs per mm gill filament: median, IQR).

### Microbiome analysis

A total of 260 host-associated SBT microbiome samples were processed through the pipeline from 65 unique fish across four body sites including gill, skin, digesta, and anterior kidney. In addition, 21 positive controls were included to determine the limit of detection of the pipeline. After applying the Katharoseq formula, the limit of detection of the pipeline, where 50% of the reads of a positive control map to the known control, was 168 reads. When the 90% threshold is applied, as recommended by the Katharoseq method, the limit of detection was 405 reads indicating that any samples with at least 405 reads could be included (Supp Figure S1). To be conservative on read depth, we included samples with at least 1000 reads and thus rarified to 1000 reads. At this sequencing depth, a total of 98 samples out of the 260 passed QC (Table 1). The majority (93.8%) of anterior kidney samples failed and thus were excluded from downstream analyses due to low successful sample size. A final table of 94 samples with 794 unique sOTU features was annotated. Digesta samples had the highest success rate (38 out of 65) followed by skin (33 out of 65) and gill (23 out of 65).

**Table 1.**
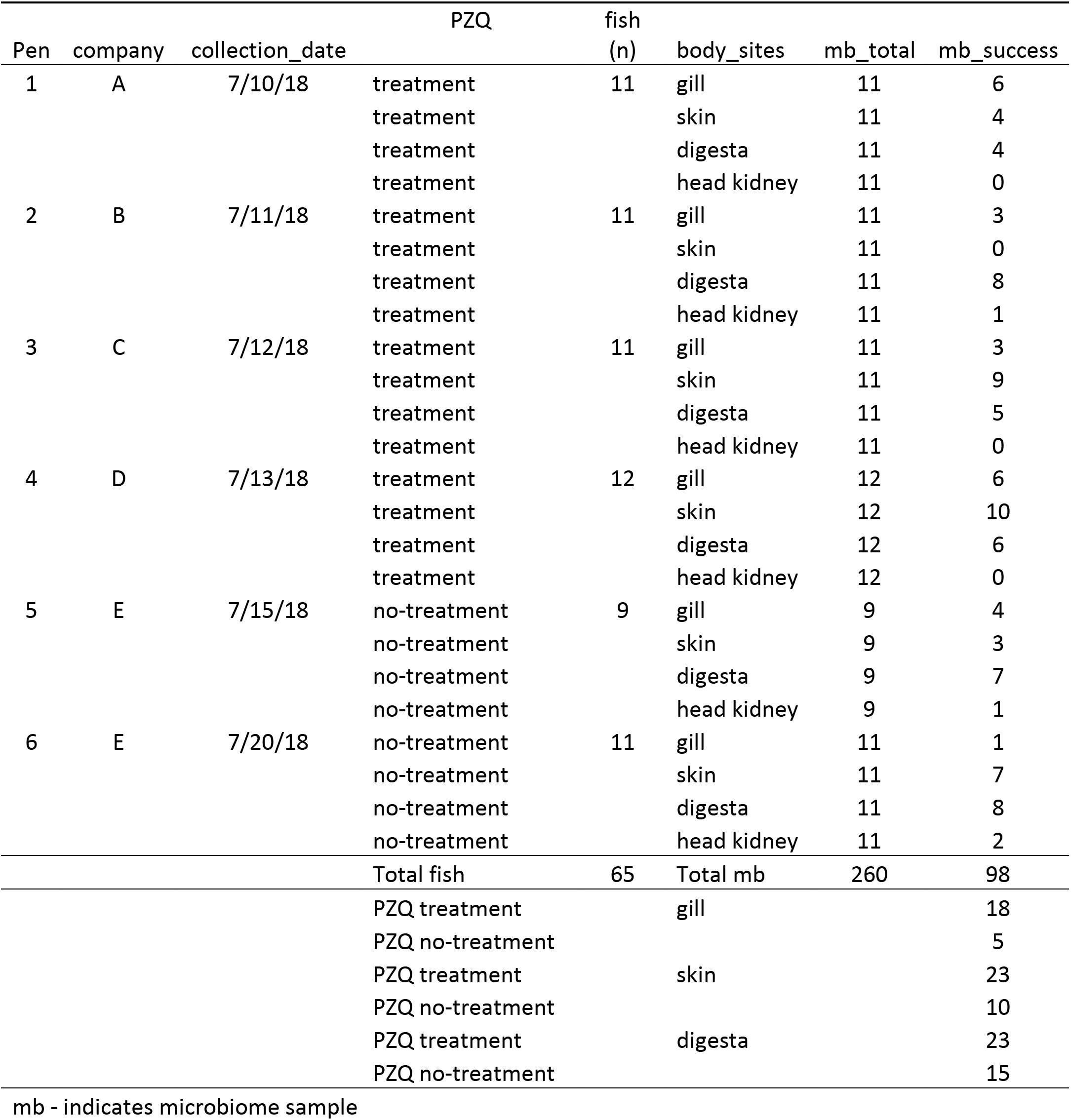
Experimental design: microbiome sampling success of Southern Bluefin Tuna

| Pen | company | collection_date | PZQ | fish<br>(n) | body_sites | mb_total | mb_success |
| --- | --- | --- | --- | --- | --- | --- | --- |
| 1 | A | 7/10/18 | treatment | 11 | gill | 11 | 6 |
|  |  |  | treatment |  | skin | 11 | 4 |
|  |  |  | treatment |  | digesta | 11 | 4 |
|  |  |  | treatment |  | head kidney | 11 | 0 |
| 2 | B | 7/11/18 | treatment | 11 | gill | 11 | 3 |
|  |  |  | treatment |  | skin | 11 | 0 |
|  |  |  | treatment |  | digesta | 11 | 8 |
|  |  |  | treatment |  | head kidney | 11 | 1 |
| 3 | C | 7/12/18 | treatment | 11 | gill | 11 | 3 |
|  |  |  | treatment |  | skin | 11 | 9 |
|  |  |  | treatment |  | digesta | 11 | 5 |
|  |  |  | treatment |  | head kidney | 11 | 0 |
| 4 | D | 7/13/18 | treatment | 12 | gill | 12 | 6 |
|  |  |  | treatment |  | skin | 12 | 10 |
|  |  |  | treatment |  | digesta | 12 | 6 |
|  |  |  | treatment |  | head kidney | 12 | 0 |
| 5 | E | 7/15/18 | no-treatment | 9 | gill | 9 | 4 |
|  |  |  | no-treatment |  | skin | 9 | 3 |
|  |  |  | no-treatment |  | digesta | 9 | 7 |
|  |  |  | no-treatment |  | head kidney | 9 | 1 |
| 6 | E | 7/20/18 | no-treatment | 11 | gill | 11 | 1 |
|  |  |  | no-treatment |  | skin | 11 | 7 |
|  |  |  | no-treatment |  | digesta | 11 | 8 |
|  |  |  | no-treatment |  | head kidney | 11 | 2 |
|  |  |  | Total fish | 65 | Total mb | 260 | 98 |
|  |  |  | PZQ treatment |  | gill |  | 18 |
|  |  |  | PZQ no-treatment |  |  |  | 5 |
|  |  |  | PZQ treatment |  | skin |  | 23 |
|  |  |  | PZQ no-treatment |  |  |  | 10 |
|  |  |  | PZQ treatment |  | digesta |  | 23 |
|  |  |  | PZQ no-treatment |  |  |  | 15 |

mb - indicates microbiome sample

**Table 2.**
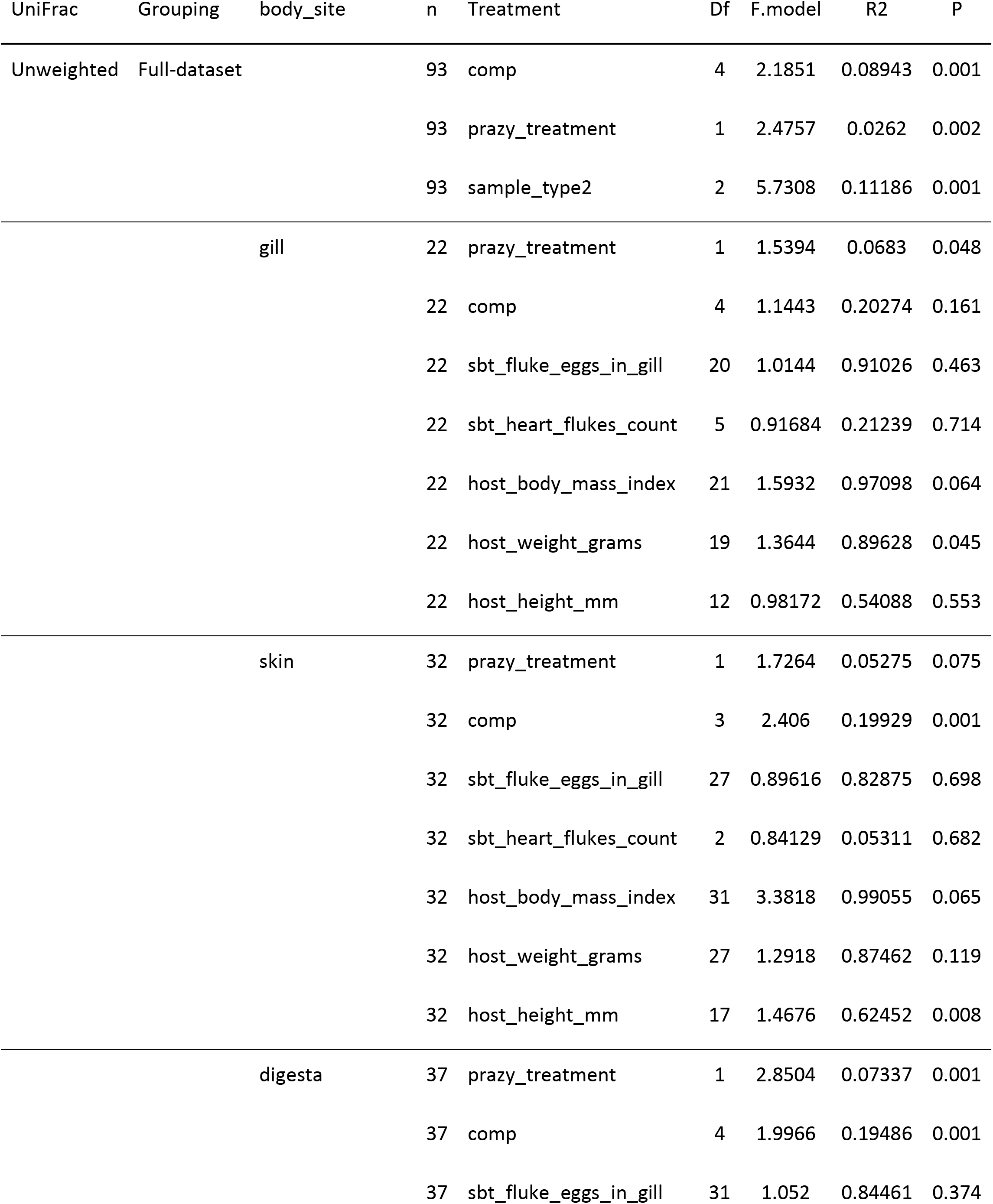
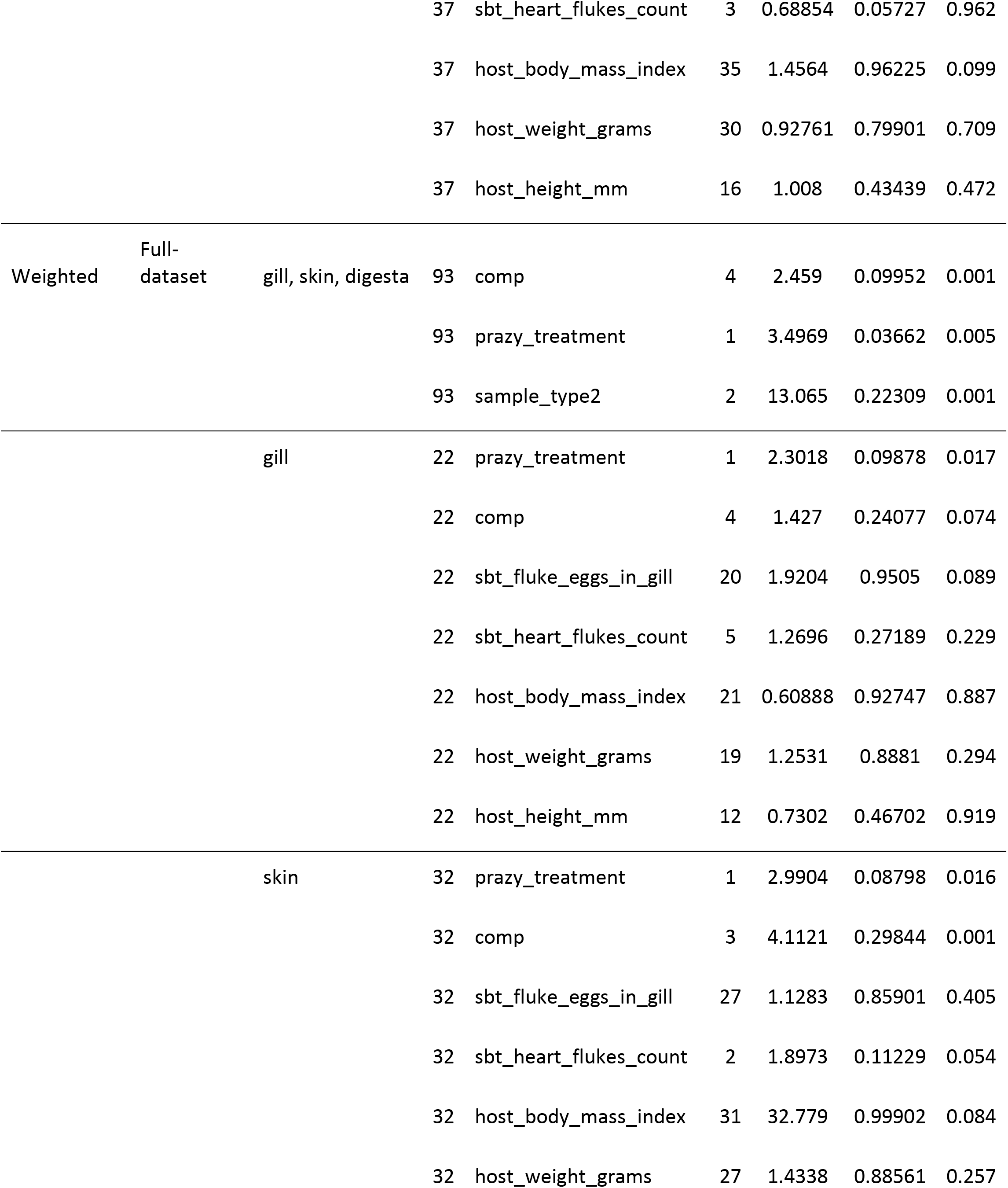
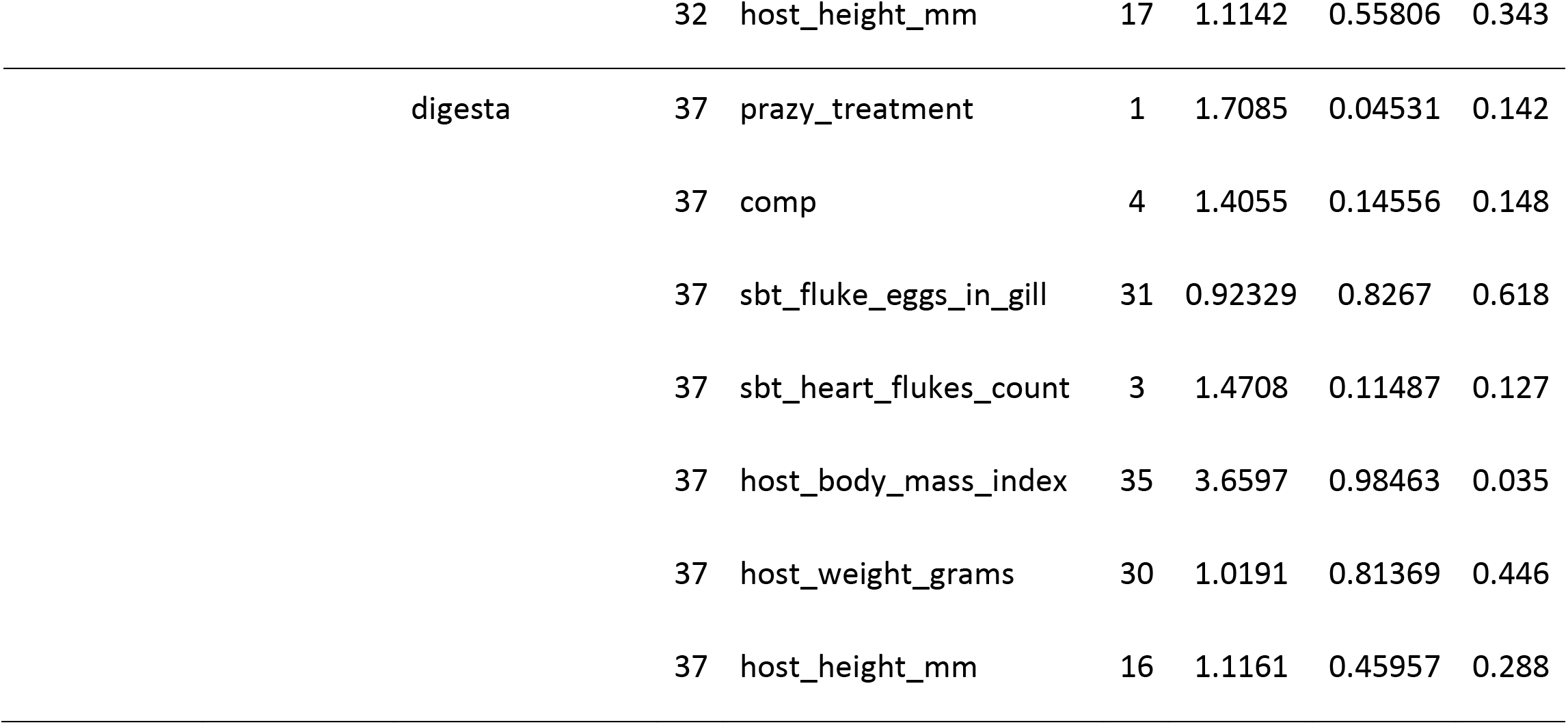
Beta diversity associations with tuna rearing condition or phenotype (adonis 999 permutations)

The microbiomes of the three body sites were compared across the fish. For alpha diversity measures, richness differed across body sites (P=0.0243, KW=7.435) with gill and skin communities having a higher richness than digesta (P<0.05) (Figure 2a). Shannon diversity differences across body sites was even stronger (P<0.0001, KW=29.97) with both gill and skin having higher evenness than digesta samples (P<0.0001) (Figure 2b). However, phylogenetic diversity did not differ across body sites (Figure 2c). Upon comparing beta diversity, samples were most strongly influenced by body site location for both Unweighted UniFrac (Figure 2d) (Adonis: P=0.001, F=5.731, R2=0.112) and Weighted UniFrac (Figure 2e) (P=0.001, F=13.065, R2=0.223) (Table 1). Microbes sampled from the SBT mucosal sites represented 20 different phyla, with digesta being enriched in Tenericutes and Spirochaetes relative to other phyla (Figure 2f). Skin and gill microbes were enriched in Proteobacteria, Cyanobacteria, Firmicutes, and Bacteroidetes relative to other phyla (Figure 2f). Microbiome variation among body sites was strongest with Weighted UniFrac. Company (geographic location) and PZQ treatment was also significant but less so than body site variation. Subsequent analyses were therefore performed on each body site independently.

**Figure 2.**
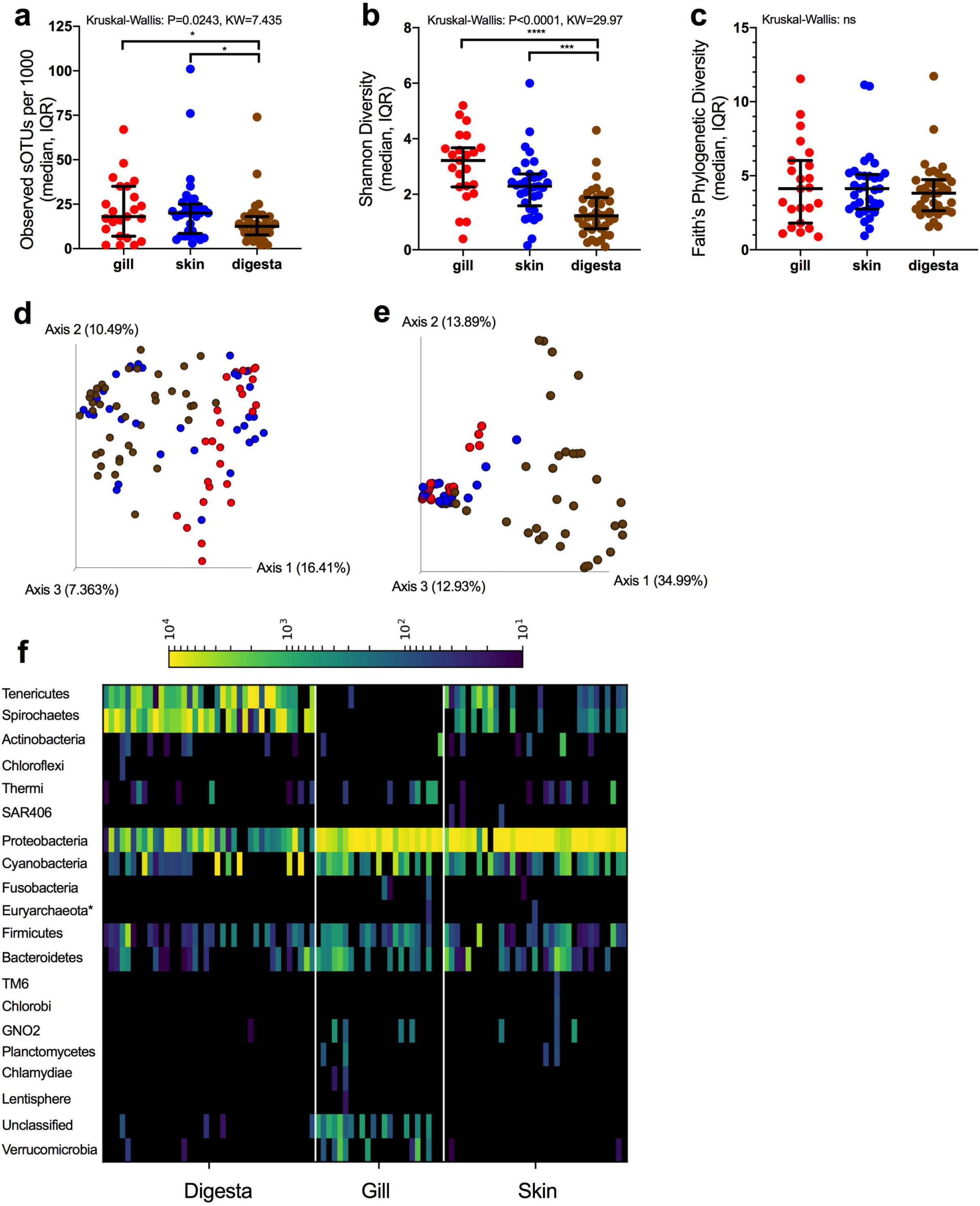
16S rRNA microbial diversity across mucosal sites in SBT. Alpha diversity measures across mucosal sites reported for a) richness, b) Shannon, and c) Faith’s Phylogenetic Diversity (median, IQR) [Kruskal-Wallis grouped test with test statistic]. Beta diversity measures of SBT colored by body site (red-gill, blue-skin, brown-digesta): d) Unweighted UniFrac and e) Weighted Unifrac. f) Phylum-level summary organized across body sites and normalized to 10,000 reads for heatmap visualization. (P<0.05 *, P<0.01 **, P<0.001 ***, P<0.0001****)

### Praziquantel treatment and the microbiome

The impact of PZQ treatment on the microbiome was evaluated using alpha and beta diversity measures. Treatment with PZQ resulted in digesta samples having lower microbial richness (Mann-Whitney: P=0.025, U=98) (Figure 3a), Shannon evenness (Mann-Whitney: P=0.0328, U=101) (Figure 3b), and phylogenetic diversity (Mann-Whitney: P=0.003, U=76) (Figure 3c). Although not significant, likely due to low sample size, all measures of microbial diversity in the gill were lower when fish were treated with PZQ.

**Figure 3.**
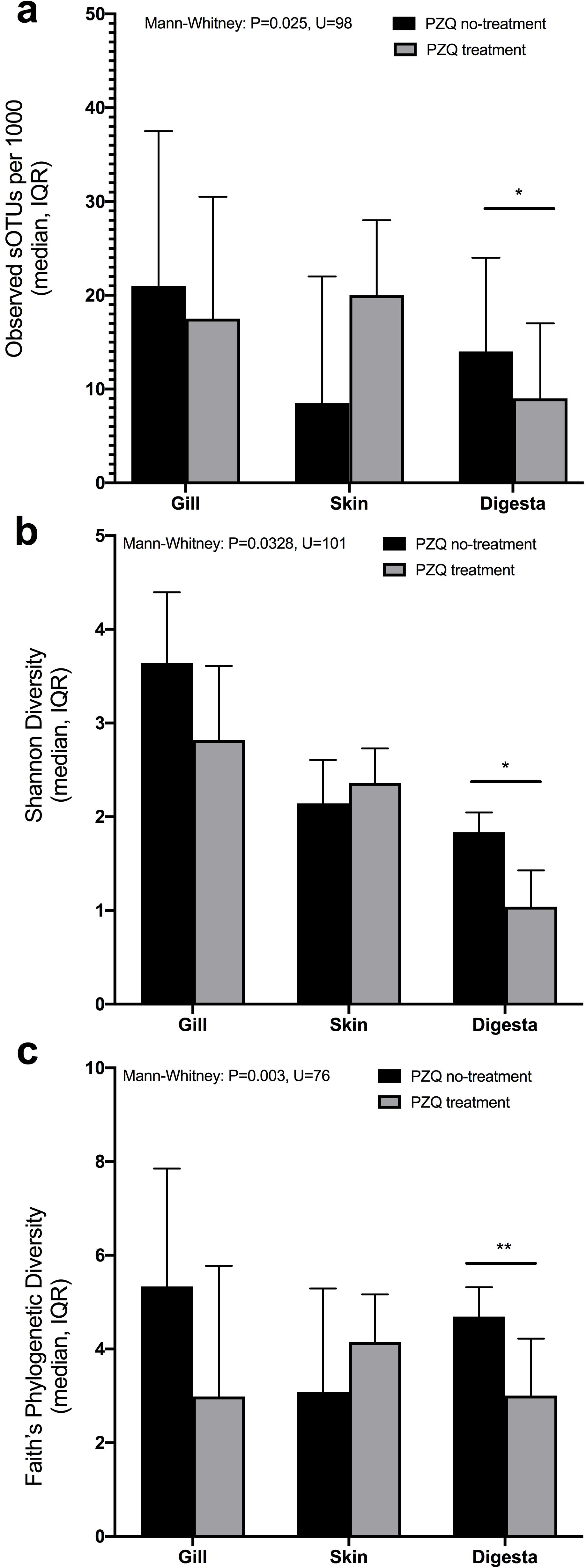
Association of PZQ treatment on alpha diversity measures across mucosal sites: a) richness, b) Shannon, and c) Faith’s Phylogenetic diversity. (P<0.05 *, P<0.01 **, P<0.001 ***, P<0.0001****)

### Cross species microbiome comparison

To understand the SBT microbiome in relation to other fish species, we performed a metanalysis comparing the SBT mucosal microbiome to a geographically (Australia) and trophically similar (tertiary carnivore) fish species, yellowtail kingfish (YTK), *Seriola lalandi* (also farmed), and a phylogenetically similar yet geographically and trophically dissimilar fish Pacific chub mackerel (MKL), *Scomber japonicus* (from the wild). Both MKL and SBT are within the same family, *Scrombridae* (tunas, mackerel, and bonito). The gill, skin, and digesta samples were similarly sampled across all species by the same researcher and further processed using the same molecular methods. Wild *S. japonicus* were sampled from the Eastern Pacific Ocean in San Diego CA as part of a fish microbiome time series study (Minich et al., 2019a). To verify that 1000 reads was sufficient to interpret and make conclusions from the data, we also compared samples rarified at 5,000 and 10,000 reads. Higher sampling depth resulted in less samples being compared, but the trends remain the same indicating that 1000 reads is sufficient for comparisons while enabling the highest number of samples to be compared (Supp Figure S2). Body sites were independently compared across all samples for both alpha and beta diversity using richness, Faith’s Phylogenetic Diversity, Unweighted UniFrac and Weighted UniFrac (Figure 4).

**Figure 4.**
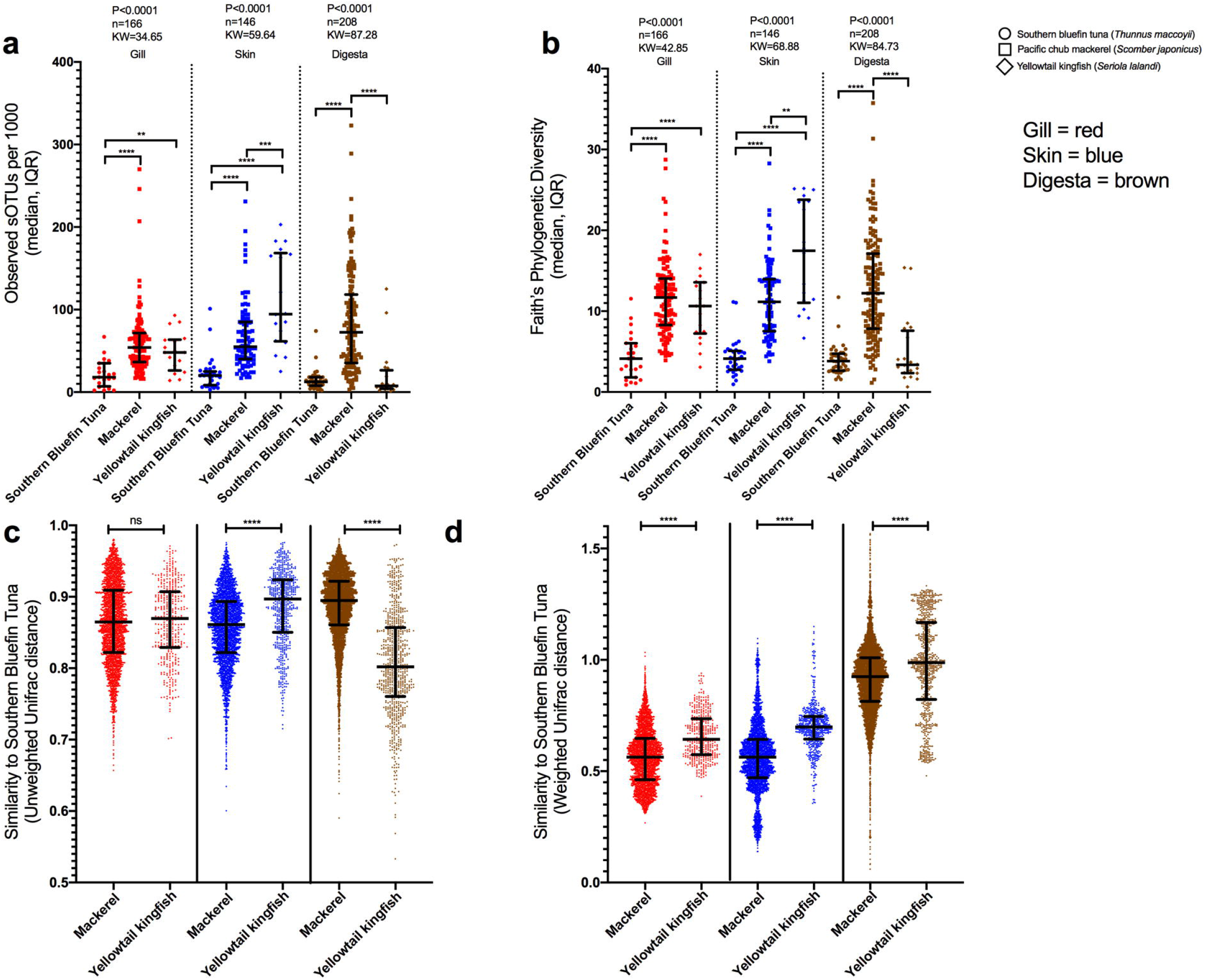
Cross fish species comparison of microbial diversity (rarified to 1000 reads). a) richness and b) Faith’s Phylogenetic Diversity across SBT, MKL, and YLK for three body sites: gill, skin, and digesta. Beta diversity comparisons of fish species MKL and YLK to SBT. Pairwise comparisons of dissimilarity from MKL and YLK each compared to SBT independent per body site using c) Unweighted UniFrac distances and d) Weighted UniFrac distances. (P<0.05 *, P<0.01 **, P<0.001 ***, P<0.0001****)

Gill microbial richness differed across species (P<0.0001, KW=34.65, n=166) with richness being lowest in SBT compared to MKL (P<0.0001) and YTK (P<0.01) (Figure 4a). Gill phylogenetic diversity exhibited the same pattern (P<0.0001, KW=42.85, n=166) with SBT having lower diversity than MKL (P<0.0001) and YTK (P<0.0001) (Figure 4b). Skin microbial diversity was also significantly different across species for richness (P<0.0001, KW=59.64, n=146) and phylogenetic diversity (P<0.0001, KW=68.88, n=146), even more so than gill microbial diversity (Figure 4a,b). On the skin, for both richness and phylogenetic diversity, a gradient was observed with SBT being lowest followed by MKL and then YTK. Digesta richness and phylogenetic diversity was lower in SBT compared to MKL (P<0.0001) while MKL was also lower than YTK (P<0.001) (Figure 4a). For phylogenetic diversity, SBT was lower than MKL (P<0.0001) for all body sites and lower than YTK for both gill and skin but not digesta (P<0.0001) (Figure 4b).

Beta diversity was assessed using Unweighted UniFrac, which gives rare taxa an equal weight, and Weighted UniFrac, which weights taxa based on their relative abundance in the sample. For each unique body site (gill, skin, and digesta), all SBT samples were compared to MKL and YTK samples rarified at 1000 reads. For Unweighted UniFrac, there was no significant difference in gill diversity (Figure 4c) whereas for skin, MKL was more similar to SBT than YTK was to SBT skin (Mann-Whitney P<0.0001, U=633804). Digesta samples of YTK were more similar to SBT as compared to MKL (Mann-Whitney P<0.0001, U=789291) (Figure 4c). For Weighted UniFrac distance, all three body sites of MKL were more similar to SBT than YTK was to SBT (gill: Mann-Whitney P<0.0001, U=341508, difference=0.080; skin: Mann-Whitney P<0.0001, U=395081, difference=0.135; digesta: Mann-Whitney P<0.0001, U=1704289, difference=0.063) (Figure 4d).

## Discussion

Parasite infections cause significant economic complications in marine aquaculture systems, particularly in high value species such as tuna (Shinn et al., 2015). In this study we set out to describe how the fish mucosal microbiome is associated by parasitic infection and treatment with praziquantel (PZQ) across three body sites including the gill, skin, and digesta of Southern Bluefin Tuna (SBT). Results indicate that the microbiome composition is most explained by body site location, followed by geographic location (company) and lastly PZQ treatment. The microbiomes at each body site were associated with company and PZQ treatment, but were not associated with blood fluke infection (measured as the number of adult blood flukes in the heart, or parasitic egg density in the gills). Lastly, in a meta-analysis, we show that mackerel may have a more similar microbial community to tuna. Since both can be infected by the same parasite and also have a similar microbiome, we suggest the feasibility of using a mackerel as a model for SBT for understanding mucosal microbiome community dynamics. Further experiments would be needed to validate the model. Our findings indicate that husbandry practices, including geographic location, influence the mucosal microbiomes of ranched SBT, and that parasite infection may not have a significant impact on fish microbiomes.

### Microbiome composition of body sites

Our overall DNA sequencing success rate was rather low for a typical microbiome study (gill = 35.4%, skin = 50.8%, digesta = 58.5%) and we attribute this to either poor field preservation or that the samples were too low of biomass. In the field, we used 95% ethanol to preserve samples, and although 95% ethanol was shown to best preserve the microbiome community for field collection in human stool samples, it is also known to inhibit various molecular assays including DNA extraction and PCR, and we would instead recommend further optimization of this preservation method in the future for fish or instead use cryopreservation, e.g. dry ice (Kemp et al., 2006; Demeke and Jenkins, 2010; Song et al., 2016). A second explanation is that the samples were low biomass meaning there was very few microbial cells. Since we used column cleanups, it’s likely that our limit of detection was between 50,000-100,000 cells, thus if a sample had less than this it would potentially fail sequencing. To improve this, we recommend using magnetic bead cleanup methods in the future (Minich et al., 2018b).

Sample type was the strongest predictor of the microbiome in the dataset. Mucosal environments of fish, including the gill, skin, and gut, are inhabited by commensal microbial communities which can have both negative and positive impacts on fish health (Gomez et al., 2013; Beck and Peatman, 2015). Recent advances in genomics methods including transcriptomics, microbiome, and proteomics have enabled the characterization and monitoring of these communities in relation to host response and environmental changes (Salinas and Magadán, 2017). Although the anterior kidney is an important organ for immune function (Abelli et al., 1994; Fänge, 1994; Watts et al., 2003; Geven and Klaren, 2017), only four of the 65 samples had detectable microbial DNA. We would expect anterior kidneys to generally lack a microbial community except in cases of systemic infection. In comparison, gill, skin, and digesta samples had a much higher success rate, indicating a rich microbial community. While various studies have focused on fish body site microbiomes independently (primarily gut followed by skin and gill), few have evaluated the cumulative microbiome across multiple body sites for individual fish (Ghanbari et al., 2015; Larsen et al., 2015; Pratte et al., 2018; Minich et al., 2019a, 2020). Our study highlights how amongst SBT body sites, gills have a high microbial diversity which was not as pronounced in the other fish species (relative to body sites within that species). Gill microbial communities in fish are understudied but have been shown to be influenced by host diet, (Pratte et al., 2018) and environmental conditions such as suspended sediment (Hess et al., 2015). Gill and skin communities however, were generally more similar and stable as compared to the gut communities in SBT. This stability has also been shown in Pacific chub mackerel, *Scomber japonicus* (Minich et al., 2019a). Gill and skin communities were primarily enriched in Proteobacteria, while gut communities had higher proportions of Tenericutes and Spirochaetes. Fish mucosal environments including the gill, skin, and digesta can also be influenced by the rearing condition and surrounding environment including the water (Minich et al., 2020).

### Influence of location or company on mucosal microbiome

When analyzing body sites independently, skin and digesta samples were significantly differentiated by the location of rearing (company). While it is probable that the five companies sampled utilized varying animal husbandry approaches (including diet) which could influence the host-associated microbial communities, it is equally likely that the pontoon location had an effect. Depth of ocean floor has been shown to positively reduce parasite infection rates in SBT as the sediment and intermediate host, the polychaete *Longicarpus modestus*, is further away from the fish and current velocities are higher (Cribb et al., 2011; Kirchhoff et al., 2011). Sediment is known to impact microbial communities of the water column, so it is likely this would additionally impact the fish microbiome (Hess et al., 2015). Another possible explanation for a company effect would be that biofouling on farming structures could differ based on location in the ocean and net cleaning frequency. Although it is not known if elevated biofouling leads to increased microbial diversity in fish, it is possible that exposure to different microbes could be greater. During pontoon aquaculture, many benthic organisms at the larval stage will settle on the pen infrastructure, utilizing nutrients from fish feces and excess feed. Many of these organisms are common benthic invertebrates and have been shown to negatively impact aquaculture by consuming oxygen, reducing water flow in the pens, and harboring microbes which can be pathogenic (Cronin et al., 1999; Fitridge et al., 2012; Madin and Ching, 2015). Biofouling communities may change depending on geospatial location of the pontoons which may experience varying oceanographic conditions including currents, wind, and upwelling.

Our study identified *Pseudoalteromonas*, *Psychrobacter*, and *Vibrio* as representing the most abundant genera across the body sites. In culture based studies, *Vibrio*, *Photobacterium*, *Pseudoaltermonas*, *Tenacibaculum*, and *Flavobacteria* have all been isolated from the gills of SBT (Valdenegro-Vega et al., 2013). Prominent SBT skin microbial genera identified included *Pseudoalteromonas*, *Vibrio*, *Acinetobacter*, *Psychrobacter*, and *Mycoplasmataceae*. Skin microbes including *Vibrio*, *Clostridium*, *Enterobacter*, *Klebsiella*, and *Proteus* isolated from other tunas have previously been shown to be important for causing decomposition through histamine production (Yoshinaga and Frank, 1982).

### Praziquantel impacts on the SBT microbiome

Treatment of PZQ had a moderate effect on microbial communities in SBT. When comparing alpha diversity, digesta samples decreased in richness, Shannon, and Faith’s phylogenetic diversity in fish treated with PZQ. When comparing beta diversity, which compares all microbes together, Unweighted Unifrac distances in gill and digesta sample community composition were influenced by PZQ treatment which indicates that rare taxa were more influenced by PZQ. For Weighted Unifrac, only gill and skin samples were influenced, indicating that more abundant microbial taxa were impacted by PZQ treatment. Humans successfully cured of schistosomiasis with PZQ had higher Fusobacterium bacteria abundances in the gut microbiome (Schneeberger et al., 2018). The associations were not strong compared to other factors, but suggests that PZQ has an impact on the human gut microbiome which could be extrapolated to fish. A caveat of this study is that only one company did not use PZQ thus the effect seen could also be confounded by husbandry differences between these companies. It is also possible that microbes in the gut may metabolize PZQ rendering it less effective or even toxic to the host (Vázquez-Baeza et al., 2018).

### Parasite – microbiome interactome

To the best of our knowledge, this is the first study to investigate the impact of blood fluke infection on the mucosal microbiome of a marine fish. However, no significant associations were observed between the microbiome and blood fluke prevalence or intensity in SBT. Hypothetically, parasitic infections can influence the microbial communities of their hosts either directly through grazing or indirectly by promoting secondary infections post trauma. In Atlantic salmon, parasitic copepod sea lice, *Lepeophtheirus salmonis*, infections were associated with decreased bacterial richness and dysbiosis of the skin microbiome (Llewellyn et al., 2017). In freshwater barramundi farms, parasitic ciliate abundance was associated with changes in the gill microbiome and mortalities (Bastos Gomes et al., 2019). Despite a relatively low incidence of infection in this study, the results suggest that blood fluke infection does not impact the microbiome. Initial studies on blood fluke densities in ranched SBT show that infected fish have on average 27 flukes per fish (Aiken et al., 2006). The median in this study was one blood fluke per fish with a maximum of six, therefore the infection intensity was low compared to previous reports.

### Modeling future SBT mucosal microbiome and parasitome resesarch

Our study demonstrated that Pacific chub mackerel (MKL), *Scomber japonicus*, has a similar gill, skin, and digesta mucosal microbiome to SBT and thus is a putative candidate model organism to study the parasitome and mucosal microbiome. Previous studies have shown that platyhelminth parasites are distributed across multiple tuna hosts (Aiken et al., 2007). Further, *Caridcola* infections occur in other Scombrid species along with other fish families (Nolan et al., 2014). *Scomber japonicus* has been shown to be successfully treated for skin fluke infections with PZQ (Yamamoto et al., 2011). For microbiome comparisons, digesta samples analyzed by Unweighted UniFrac and alpha measures showed yellowtail kingfish (YTK) was more similar to SBT. Since YTK and SBT are both tertiary carnivores and, in this study, both are farmed, their diet is generally more rich in fish protein than MKL which are small pelagic secondary consumers (Agusa et al., 2007; Hajeb et al., 2010). This association of diet and high trophic level corresponding with low diversity was first observed in mammals, and may also be conserved to some extent in fish (Ley et al., 2008a, 2008b). However, when relative abundance (compositional) measurements are included in the analysis, all MKL body sites were more similar to SBT, which could suggest evidence of phylosymbiosis for highly abundant taxa. For microbiome studies, a link has been demonstrated within plants and animals to suggest that phylogenetically related hosts retain more similar microbiomes (Pollock et al., 2018; Ross et al., 2019; Lim and Bordenstein, 2020). While most of these studies have focused on the gut microbiome, additional body sites have recapitulated this relationship in the skin (Chiarello et al., 2018) but not gills for fish and amphibians. Our study demonstrates how gill, skin, and gut communities of the SBT were more similar to a genetically similar fish within the same family.

Since the MKL microbiome is similar to SBT, it provides an opportunity to explore developing MKL as a model for future SBT research. One of the challenges with marine finfish aquaculture is performing replicated experiments, including disease challenges, that are not feasible to do in a production setting (Salama and Rabe, 2013). Mackerel is an ideal species for aquaculture research as they are globally distributed, not threatened, easy to catch, inexpensive, small, and easy to culture in tanks.

## Conclusion

Our findings demonstrate that Southern Bluefin Tuna harbor a unique microbiome in each body site. The SBT microbiome is influenced by pontoon location and, to a lesser extent, treatment with the antihelminth therapeutic, praziquantel. No association between blood fluke infection and the microbiome was identified. It is possible, however, that infection intensity was not sufficient or sample size not adequate to identify a relationship. In addition, we showed genetically similar fish (Scombrids) have a more similar microbial community across multiple body sites including gill, skin, and digesta, suggesting possible phylosymbiosis. Based on this finding, the exploration of Pacific chub mackerel as a candidate model organism for studying the microbiome and potential parasitome of SBT is proposed. Tuna are highly valued, and present significant challenges for conducting controlled experiments due to their size and physiology. Future work employing a smaller, highly accessible relative could enable greater gains in research and ultimately, enhanced aquaculture productivity.

## Permission to reuse and Copyright

We give permission to published all Tables and Figures which were generated by the authors.

## Data availability Statement

Data is made publicly available on Qiita (12227), Qiita analysis (28067). Sequence data is also submitted to EBI (ERP120036).

## Ethics Statement

Microbiome samples were collected during a harvest event on dead fish in coordination with the companies according to standards outlined by The Australian Southern Bluefin Tuna Industry Association.

## Author Contributions

JJM, BN, NJB, and EEA contributed to the conception and design of the study. JJM collected microbiome samples in the field, performed DNA extractions and microbiome processing, analyzed the data and wrote the first draft of the manuscript. CP performed analyses on gill egg density and heart fluke counts. MM contributed to analysis and visualization. All authors contributed to the manuscript revision, and read and approved the submitted version.

## Acknowledgements

We thank the University of Tasmania for hosting J.J.M. and Royal Melbourne Institute of Technology (RMIT) for enabling sample processing. The Australian Southern Bluefin Tuna Industry Association are acknowledged for hosting the team in Port Lincoln, providing support for the collection of samples, including visits to commercial SBT ranching pontoons.

