## Supplemental Figures for "The Southern Bluefin Tuna mucosal microbiome is influenced by husbandry method, net pen location, and anti-parasite treatment"

*Supplementary Material*

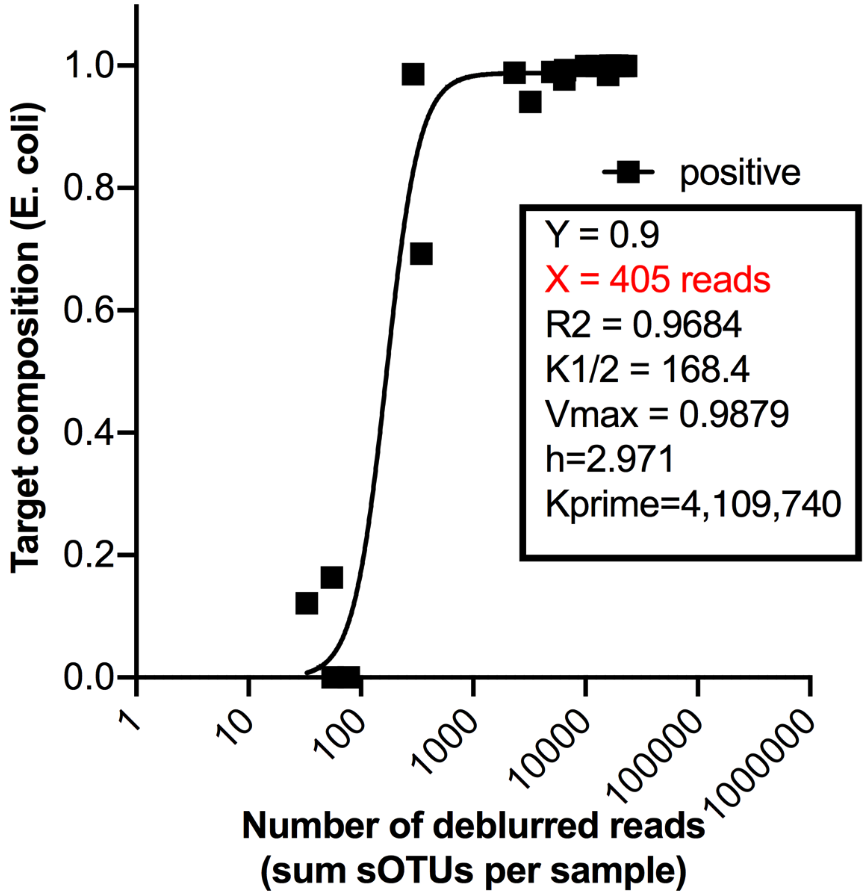


**Supplementary Figure 1.** Sample exclusion cut off determined by comparing relative abundaces of expected target sequences across a serial dilution of positive controls. See Katharoseq method for more details.

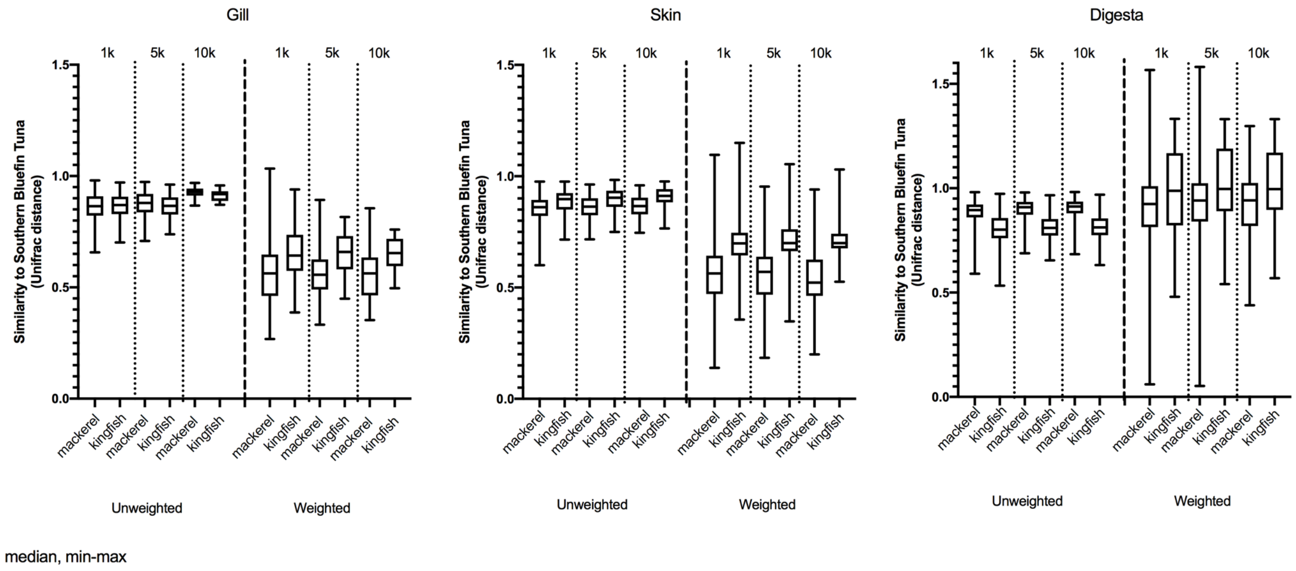


**Supplementary Figure 2.** Justification for setting a rarefaction depth of 1000 reads for cross-species comparisons of microbiome. Samples of gill, skin, and digesta are uniquely compared between fish species: mackerel and yellowtail kingfish to Southern Bluefin Tuna for Unweighted and Weighted Unifrac distances.
